# *Drosophila SA1* expression prevents brain tumorigenesis and PARP-mediated cell elimination

**DOI:** 10.1101/2025.04.18.649500

**Authors:** Simona Totaro, Antonella Lettieri, Silvia Castiglioni, Francesco Lavezzari, Cristina Gervasini, Valentina Massa, Thomas Vaccari

**Affiliations:** Department of Biosciences, Università degli Studi di Milano, Milano, Italy; Department of Health Sciences, Università degli Studi di Milano, Milan, Italy

## Abstract

The cohesin complex performs essential cellular functions including regulation of chromosome cohesion, chromatin organization and DNA repair. Somatic pathogenetic variants in cohesin genes, such as *STAG2*, have been associated with cancer, but their contribution to brain tumorigenesis is unclear. Here, we report the presence of *STAG2* variants in glioblastoma and medulloblastoma patients and determine that loss of *STAG2* in human cells leads to DNA damage and apoptosis. Treatment with inhibitors of the Poly ADP-ribose polymerase (PARP), which are used to treat forms of cancer with defects in DNA repair, increased the amount of apoptosis, confirming that synthetic lethality between reduced cohesin and PARP activity could be observed *in vitro*. Similar results were obtained *in vivo* by reducing expression of *SA1,* the *Drosophila melanogaster* homolog of *STAG1/2.* Cohesin gene silencing during fly brain development leads to defects in neural stem cells differentiation and tumorigenesis both in the presence of oncogenic activity and *per se*. Our *in vivo* and *in vitro* data suggests that impairment of PARP activity might induce synthetic lethality in cohesin-dependent tumors, highlighting a vulnerability that can be pharmacologically exploited.

## INTRODUCTION

Cohesin proteins are part of a conserved ring complex that ensures sister chromatid cohesion (Nasmyth, 2011). The cohesin complex also regulates genomic stability by taking part in DNA repair. In addition, it orchestrates gene expression by acting on the genome 3D architecture and therefore, participating in chromatin remodeling (Nasmyth and Haering, 2009). While cohesin genes are essential for survival, heterozygous germline loss of function variants cause congenital disorders called cohesinopathies, including Cornelia de Lange and Roberts syndromes (Kline *et al*., 2018).

In humans, the cohesin core complex is formed by SMC1A, SMC3, RAD21 and STAG1, STAG2 or STAG3 (Nasmyth and Haering, 2009). While STAG3 is essential for proper chromosome pairing and segregation in meiosis (Bayés *et al*., 2001), STAG1 and STAG2 have broader, partially overlapping functions. However, STAG2 is specifically required for regulation of transcription in DNA repair (Romero-Pérez *et al*., 2019).

Owing to the many tumor suppressive processes in which the cohesin complex is involved, somatic mutations in multiple cohesin genes were found in different types of tumors (Di Nardo, Pallotta and Musio, 2022). In particular, *STAG2* is a frequent target of inactivating mutations in human cancers (Hill, Kim and Waldman, 2016; Arruda *et al*., 2020). These usually include frameshift, nonsense, or splice site mutations leading to aberrant proteins (De Koninck and Losada, 2016). *STAG2* variants were identified in a variety of tumors, including glioblastoma, (Solomon *et al*., 2011), urothelial bladder cancer (Balbás-Martínez *et al*., 2013; Guo *et al*., 2013; Solomon *et al*., 2013; Taylor *et al*., 2014; Tirode *et al*., 2014; Weinstein *et al*., 2014), melanoma (Solomon *et al*., 2011) myelodysplastic syndrome, acute myeloid leukemia (Solomon *et al*., 2011) and Ewing’s sarcoma (Brohl *et al*., 2014; Crompton *et al*., 2014). Somatic variants in *STAG1* are also involved in the tumorigenesis of colorectal cancer, bladder cancer, Ewing’s sarcoma and myeloid malignancies (Balbás-Martínez *et al*., 2013; Kon *et al*., 2013; Thota *et al*., 2014; Weinstein *et al*., 2014).

Cohesin genes are functionally conserved in *Drosophila melanogaster.* In particular, the fly genome encodes two homologs of human *STAG1-3*. *Stromalin1* (*SA1 or* CG3423) is ubiquitously expressed and appears to be related to *STAG1-2*, while *SA2* (CG13916) is mainly expressed in male gonads and is likely functionally similar to *STAG3* (Thomas *et al*., 2005)*. SA1* was initially identified with other fly cohesin genes for its ability to bind chromatin and regulate enhancer-promoter communication and support sister chromatid cohesion (Rollins *et al*., 2004; Gause *et al*., 2008). During embryogenesis, *SA1* expression is controlled by Notch target Cut in neuroblasts (NB). *SA1* expression in such context supports developmentally regulated NB death, preventing the emergence of ectopic NBs. Such process appears to be regulated cohesin ability to regulate chromatin architecture (Arya *et al*., 2019). In the developing larval fly brain, *SA1* has also been implicated in post-mitotic regulation of morphogenesis, in particular neuronal pruning, a process dependent on transcriptional regulation (Schuldiner *et al*., 2008), as well as in establishing the pool of synaptic vesicles for memory formation (Phan *et al*., 2019). A number of genetic manipulations in *Drosophila* have been previously used to assess the contribution of specific genes to differentiation of neural stem cells and to mimic brain tumorigenesis (Read *et al*., 2009; Paglia *et al*., 2017; Maurange, 2020).

The poly ADP-ribose polymerase (PARP) is a central sensor of DNA damage. Drugs that inhibit PARP activity are currently used to induce synthetic lethality of breast and ovarian cancer cells with mutations in genes regulating DNA repair pathways (D’Andrea, 2018; Mekonnen, Yang and Shin, 2022). A few lines of evidence in *in vitro* systems have reported that depletion of *STAG2* causes susceptibility to PARP inhibitors (Mondal *et al*., 2019; Luo *et al*., 2024). While the first evidence of *STAG2* involvement in tumorigenesis was the presence of focal deletions on the X chromosome in glioblastoma (Solomon *et al*., 2011), no animal models of brain tumorigenesis based on reduced *STAG1/2* in somatic cells exist and no study of the effect of PARP inhibitors *in vivo* have been reported.

Here, we updated the repertoire of *STAG2* variants associated with glioblastoma and medulloblastoma. We also determined the consequences of depleting *STAG2* in cells on DNA repair and apoptosis with and without PARP inhibitor supplementation. We have found that *STAG2* deficiency leads to persistent DNA damage and that treatment with a PARP inhibitor increased the amount of apoptosis in spheroids, when compared to those with normal *STAG2* expression. To model the effect of somatic *STAG2* deficiency *in vivo,* we reduced the expression of *Drosophila SA1* in developing larval neuroblasts, the brain stem cells that give rise to neurons and glia. We observed an impairment of NB differentiation during larval development that correlates with occasional formation of tumor-like masses in adult brains and, in general, to a shortened lifespan. RNA interference (RNAi) against *Drosophila Parp1,* the unique homolog of human PARP enzymes, or PARP inhibitor supplementation, does not appear to alter NB differentiation significantly. However, combined reduction of *SA1* and *Parp1* activity reverts *SA1* NB phenotypes. We also find that in epithelial tissue, reduced *SA1* and *Parp1* expressions lead to additive stabilization of DNA damage, suggesting that the amelioration of tumor-associated SA1 phenotypes might be due to synthetic lethality. Our data indicate that cohesin genes act as tumor suppressors and that their loss can be compensated by PARP inactivation *in vivo*. Thus, tumors with cohesin mutations might be highly dependent on DNA repair pathways, representing potential therapeutic targets for PARP inhibitors.

## METHODS

### Cell cultures and cell-based assays

HEK-293T cells were grown in DMEM medium (Dulbecco’s modified Eagle’s medium, Life technologies, 11965092) supplemented with 10% FBS (fetal bovine serum, Life technologies, 10500064) and 1% of penicillin-streptomycin (P/S) (Euroclone, ECB3001D). Cells were cultured in a Petri dish at 5% CO2 and 37 °C.

HEK-293T shSCR and HEK-293T shSTAG2 are derived from HEK-293T control (CTRL) cell line obtained through viral infection with a control scrambled (SCR) plasmid and the one containing a short hairpin (sh) for the STAG2 gene silencing (shSTAG2); both plasmids also contain GFP encoding gene and puromycin cassette for selection. Briefly, STAG2-shRNA lentiviral vector (Origene, Locus ID 10735) or the control scrambled (SCR) sequence SCR-shRNA as well as packaging and envelop virus components was transfected into HEK-293T cells by using CaCl_2_ method. After two days, medium containing virus particles were collected and used to infect new HEK-293T to generate stable cell lines. Cells were positively selected with puromycin (1ug/mL Invivogen ANT-PR-5) treatment for 72 hours (h) 48h post-infection, and then periodically maintained under selection.

To perform MTT assays, 20000 cells/well were seeded in a 24 multiwell till 70% of confluence. Then, the culture medium was removed, and 300µl of serum-free blank medium and 30µl of MTT stock solution (5mg/ml MTT powder Sigma-Aldrich, M2003) were added to each well. The plate was incubated at 37°C for 30min/2 hours until purple precipitates appear. After removing the solution, 300ul of isopropanol were added to each well, and the plate was shaken for 10 minutes at RT. After resuspension, 200µl of each well were transferred into a 96 multiwell with lid. The levels of precipitates were read by Insite spectrophotometer at 570nm.

For adhesion tests, cells in triplicates were seeded at a concentration of 40.000 cells per well in a 24 multiwell containing a glass slide, then the plate was incubated for 2 hours at 37°C. After medium removal, cells were fixed with 300µl of cold methanol for 10 minutes at –20°C. After fixation each well was washed twice with PBS 1X for 5 minutes and 300µl of hematoxylin were added for 1 minute at RT. After hematoxylin removal, each well was washed with water until it became clear and 300µl of eosin were added for 1 minute at RT. Each well was washed with water and slides were mounted on a slide with Mowiol.

To assess spheroid formation, 5000 cells/well were seeded in a low attachment 96 U-bottom multiwell. Cells were resuspended in DMEM/F12 medium (Gibco, 11320033) containing B-27 supplement (100X, Gibco, 12587010), N-2 supplement (50X, Gibco, 17502048), heparin (2 μg/ml, Merck Life Science, H3149-50KU), EGF (20 ng/mL, Prepotech, AF-100-15-500UG), FGF2 (10 ng/mL, Prepotech, AF-100-18B-500UG), and P/S 100X. Then, 200ul/well were aliquoted and the plate was centrifuged for 5 min at 300g. Spheroids of each cell line were grown for 4 days and then moved in a 24-multwell plate previously coated with Matrigel (Merck Life Science, CLS356234-1EA) to evaluate dissemination ability.

### Drosophila methods

*Drosophila* strains were maintained in vials containing a standard food medium composed of water, 34% cornmeal, 57% molasses, 9% yeast, 0.7% agar, 0.7% propionic acid and 2% tegosept. Supplemented food composition was modified by replacing molasses with 35% sucrose and with 44% yeast concentration to enhance egg-laying. Unless otherwise specified, fly lines were generated by crossing or recombination from stocks obtained in our laboratory or sourced from the Bloomington Drosophila Stock Center (BDSC), Indiana University, and the Vienna Drosophila Resource Center (VDRC). Crosses were kept at 25°C. The following fly stocks were used:

| Code | Origin | Genotype |
| --- | --- | --- |
| 64349 | BDSC | *CantonS* |
| 31603 | BDSC | *y[1] v[1];+; P{y[+t7.7] v[+t1.8]=TRiP.JF01355}attP2* |
|  | This Work | *w118, ms1096Bx-GAL4,P{w[+mC]=UAS-EGFP}8; +; +* |
|  | Cédric Maurange, Aix Marseille University | *UAS-dicer2;wor-GAL4, ase-GAL80;UAS-mCD8::GFP* |
| 36794 | BDSC | *y[1] sc[\*] v[1] sev[21]; P{UAS-TRiP.GL00534}attP40/CyO,Tb; +* |
| 33395 | BDSC | *y[1] sc[\*] v[1] sev[21]; +; P{y[+t7.7] v[+t1.8]=TRiP.HMS00272}attP2* |
|  | McKim, The State University of New Jersey | *+, SA[ex86] pr FRT40A / CyO; +* |
|  | This Work | *+, P{UAS-TRiP.GL00534}attP40/CyO,Tb;P{GD9445}v46745* |
|  | This Work | *+, SA[ex86] pr FRT40A/CyO,Tb;P{GD9445}v46745* |
| 46745 | VDRC | *w1118;+;P{GD9445}v46745* |
|  | Renee Read, Emory University | *UAS DP110-CAAX; +; Repo GAL4-UASmCD8GFP,UAS ΔEGFR/TM6c,Tub-GAL80* |
|  | Renee Read, Emory University | *+, +; Repo GAL4-UASmCD8GFP/TM2* |
| 65229 | BDSC | *y[1] sc[\*] v[1] sev[21];P{y[+t7.7] v[+t1.8]=TRiP.HMC06090}attP40; +* |
| 36598 | BDSC | *y[1] sc[\*] v[1] sev[21]; +; P{y[+t7.7] v[+t1.8]=TRiP.GL00558}attP2* |
| 36783 | BDSC | *y[1] sc[\*] v[1] sev[21]; P{y[+t7.7] v[+t1.8]=TRiP.GL00518}attP40; +* |
| 36794 | BDSC | *y[1] sc[\*] v[1] sev[21]; P{TRiP.GL00534}attP40 / CyO; +* |
| 41976 | BDSC | *y[1] sc[\*] v[1] sev[21]; +; P{y[+t7.7] v[+t1.8]=TRiP.HMS02374}attP2* |

For *in vivo* PARP inhibitor supplementation, 3-AB (Catalog No. S1132, Selleck USA) was used as PARP inhibitors. A 50 mM stock solution was prepared in DMSO following manufacturer’s datasheet and diluted in sterile distilled water to final concentrations of 5–250 μM (150 μL/tube). These were added to 3 mL of *Drosophila* white food, pre-punctured with 20 holes for uniform diffusion, resulting in final food concentrations of 0.25–12.5 μM. Equal DMSO volumes served as controls. After the housing period on standard food, flies were transferred to treatment (3-AB) or control (DMSO) vials and moved to fresh food every two to three days.

To score fly survival, A maximum of 20 flies/genotype (equally distributed in male and female) were kept in a single vial and kept at 25°C. In order to perform a good statistical analysis, each lifespan assay was carried out on ∼100-150 flies for genotype. For each experiment, three independent biological replicates were counted both for experimental and control progenies. Flies were transferred to fresh food medium every two to three days, when dead flies were also scored. The counts were analyzed with PRISM GraphPad software: survival fractions were calculated using the product limit Kaplan-Meier method and analyzed with the Gehan-Breslow-Wilcoxon and Long-rank (Mantel-Cox) test using Prism software to evaluate significance of differences between survivorship curves.

### Real-time quantitative PCR

For cell culture experiments, 1µg of RNA extracted with phenol/chloroform method by NucleoZOL (Macherey-Nagel) was retrotranscribed by using All-In-One 5X RT MasterMix (Microtech) kit. Quantitative real-time PCR (RT-qPCR) TB Green Premix Ex Taq (Tli RNase H Plus) (Takara) kit and the CFX Opus 96 Real-Time PCR System (Bio-Rad) were used to evaluate gene expression following manufacturer instructions. The data obtained from RT-qPCR were analyzed using a comparative Ct quantification method. ΔCt was obtained by normalizing each Ct sample to the housekeeping genes (*GAPDH, RPLP0, RPL13A*) mean. Then the ΔΔCt was obtained by comparing the ΔCt of every sample for each gene to the reference one gene expression of the treated samples against its control. Relative gene expression values were obtained by calculating the Fold change (FC) (2 –Δ ΔCT). Technical and biological triplicates were performed for all the experiments. The following primers were used for RT-qPCR: *GAPDH* (For: 5’-AGCCACATCGCTCAGACAC; Rev: 5′-GCCCAATACGACCAAATCC) *RPLP0*(For: 5’-TCTACAACCCTGAAGTGCTTGAT; Rev. 5′-CAATCTGCAGACAGACACTGG) *RPL13A* (For: 5’-CCTGGAGGAGAAGAGGAAAGAGA; Rev: 5′-TTGAGGACCTCTGTGTATTTGTCAA)

*STAG2* (For: 5’-AAGGAGGACTTGCTGCGTTT; Rev. 5′-

TCCTCTTGCTGACCATCTGC). RT-qPCR and western blot data were performed using Student t-test, considering significance for p-value < 0.05 (* p < 0.05; ** p < 0.001; *** p < 0.0001).

For *in vivo* experiments, after dissection, organs of the selected genotype were homogenized in Trizol reagent (Invitrogen, 15596-018). For wing disc 30 discs were used, while for brain 20-25 organs dissected at different stages of development were used. Timepoint analyzed were larva L3, pupa at stages P7-8 and P11-12, young adults (5 days old, equal number of male and female), adults (10 days old) and aged adults (25 days old).

The RNA extraction was performed using a commercial kit (Kit Zymo RNA extraction insect tissue). The concentration of extracted RNA and DNA were measured using the NanoDrop 1000 Spectrophotometer. Retrotranscription of RNA to cDNA was performed using LunaScript® RT SuperMix Kit (New England BioLabs, E3010). RT-qPCR was performed with Luna® Universal qPCR Master Mix (New England BioLabs, M3003) by using the CFX Connect Real Time PCR Detection System (Bio-Rad, 1855201).

Results were normalized using the housekeeping *Rp49* and the ΔΔ cycle threshold method and results expressed as the relative change (-fold) of the downregulated group over the control group, which was used as a calibrator.

*GFP*, *SA* and *Rp49* RT-qPCRs were performed using Sybergreen (Applied Biosystems) with the following primers:

*GFP* (For: 5′-GTCAGTGGAGAGGGTGAAGG; Rev: 5′-TACATAACCTTCGGGCATGG),

*SA (*F: 5’-TTGTGCGACACTCGAAGAAC; R: 5’-CCGCTTTCTTCGTCAAACTC),

*Rp49 (*F: 5′-ACGTTGTGCACCAGGAACTT; R: 5′-TACAGGCCCAAGATCGTGAA).

Output data were analyzed using CFX Manager Software (BioRad) and Prism that was used also to realize graphs.

### Protein extraction and quantification and Western blot and analysis

For cell extracts, HEK-293T pellets were resuspended in cold S300 buffer (NaCl 300mM, HEPES pH 7.6 50mM, NP40 0,1%, MgCl2 2mM, glycerol 10%) added with protease and phosphatase inhibitors and GENIUSTM Nuclease (SC-202391). Samples were left on ice for 1 hour and then centrifuged at maximum speed for 10 minutes at 4°C. Supernatants containing protein lysates were collected and quantified using Bradford method. BSA 0,2mg/ml (Bio-rad 5000206) was used to prepare the standard curve. Finally, samples were predisposed for the western blot run by preparing aliquots at 1µg/µl supplemented with Laemmli sample buffer 4x (LSB)(Bio-Rad, 1610747) and boiled for 10 minutes at 100°C. 30µg of samples were loaded and separated by SDS-PAGE (Running buffer 1x diluted from 10x made of 3% Tris HCl, 14,4% Glycine and 1% SDS) using 10% polyacrylamide gels (Biorad Mini-PROTEAN TGX Gels, 4561034). At the end of the run, cold transfer buffer 1X (20% methanol and 10% Transfer buffer 10x, composed of 3% Tris HCl and 14,4% Glycine) was used to transfer protein samples to a nitrocellulose membrane (Merck Life Science, GE10600003). Milk (Sigma-Aldrich, 4259001) 5% in TBS-T 1X was used as a blocking solution for 1h at RT, then membranes were incubated with primary antibody diluted in milk or TBS-T 1X at 4°C O/N (rabbit anti-STAG2, 1:1000, Cell Signaling, 5882; rabbit anti-γH2A.X, 1:1000 Cell signaling, BK9718S; rabbit anti-GAPDH 1:1000, Cell signaling, 5174). Then, membranes were incubated with secondary antibody anti-rabbit or anti-mouse HRP 1:3000 (Biorad, rabbit 1706515, mouse 1706516) diluted in TBS-T 1X or in milk for 1h at RT. Membranes were washed three times with TBS-T 1X and incubated with ECL or Amersham to detect chemiluminescence signals captured by Chemidoc Imaging System. Data obtained from western blots were analyzed using ImageJ software, the mean pixel intensity was calculated, and the t-student Test was used for statistical analysis. Experiments were performed in biological and technical triplicates.

### Immunofluorescence analyses

Cells were permeabilized for 10’ with PBT (PBS with 0,25% Triton) and then natural donkey serum (20% of NDS in PBT 0,25%) was used as blocking solution for 1h at RT. Cells were incubated with primary antibody rabbit anti-γH2AX (1:200, Cell signaling, BK9718S), and rabbit anti-cleaved caspase 3 (1:200, Cell signaling, BK9664S) at 4°C overnight. The day after, secondary antibodies donkey Alexa Fluor Cy-3-conjugated anti-rabbit Fab fragments (1:200; Jackson Immunoresearch) were used and incubated for 2 hours at RT. Then, cells were washed with PBT 0,25% three times and DAPI (1:1000) was used for nuclei counterstaining. Finally, cells were washed with PBS 1X and with MilliQ water. Slides were mounted with Mowiol (Sigma Aldrich), and signals were acquired by fluorescence microscope at a 10x magnification and for cCas3 quantified by ImageJ software.

For immunolabeling of *Drosophila* organs, adult brain were dissected and processed as described (Wu and Luo, 2006; Ostrovsky, Cachero and Jefferis, 2013). Larvae were reared for 120– 150 hours post-egg deposition and wandering third-instar larvae were selected for analysis. Larval brains and wing discs remained attached to carcasses for ease of handling. Carcasses were prepared by removing the gut, fat tissue, and salivary glands in 1× PBS, then fixed in 4% PFA for 20 min at room temperature. Tissues were rinsed three times in 0.1% Triton X-100 in 1× PBS (PBST) for 5 min to remove fixative, followed by permeabilization in 0.3% Triton X-100 in 1× PBS (PBST 0.3%) for 30 min. Blocking was performed with 5% BSA in PBST 0.3% for 30 min before overnight incubation with primary antibodies diluted in blocking solution. The following primary antibodies were used: chicken anti-GFP (1:1000, Abcam, 92456), rabbit anti-Miranda (1:500, Abca, 197788), mouse anti-Prospero (1:100, DSHB, 528440), rat anti-Elav (1:50, DSHB, 528218), mouse anti-histone 2A gamma variant, phosphorylated (γH2Av) (1:50, DSHB, 2618077), rabbit anti-Fibrillarin (1:500, Abcam, 5821), mouse anti-nc82 (1:40, DSHB, 2314866). After three washes, Alexa Fluor-conjugated secondary antibodies (1:300, Invitrogen) were incubated for 2 hours at room temperature. DNA was stained with DAPI (1:5000, Sigma Aldrich), followed by three washes and fine dissection. Samples were mounted in Moviol (Sigma Aldrich) and dried overnight at room temperature.

### Microscope acquisition

Brightfield images of spheroids were acquired with 4x magnification. Confocal images were acquired with a Nikon A1R/AX laser scanning confocal microscope equipped with a Nikon A1/AX plus camera and the following objectives (Nikon): Plan Fluor 10X DIC L N1 (NA 0.3), Plan Fluor 20X DIC N2 (NA 0.5); DAPI, Alexa Cy3 were excited at 405, 561 nm and observed at 425–475, 570–620 nm, respectively.

Images of *Drosophila* organs were acquired using Nikon A1-SIM or NiU confocal microscopes from the UniTECH NoLimits departmental platform (https://unitech.unimi.it/). Immunofluorescence images were analyzed using FIJI software FIJI (Fiji is just ImageJ, NIH), followed by statistical analysis with Prism (GraphPad software version 9.1.2 La Jolla, CA, USA; www.graphpad.com). Measurements and fluorescence evaluation were carried out through the FIJI Software. Quantification analysis was performed by evaluating equal numbers of Z-stack among different genotypes for comparative analysis.

### Database searches

The publicly accessible cBioPortal for cancer genomics database (https://www.cbioportal.org/) was investigated, selecting datasets from 8 studies of medulloblastoma (Jones *et al*., 2012; Pugh *et al*., 2012; Robinson *et al*., 2012; Poeran, 2016; Northcott *et al*., 2017) and glioblastoma (Brennan *et al*., 2013; Zhao *et al*., 2019; Wang *et al*., 2021). The CADD score was calculated for all SNP *STAG2* variants reported in gnomAD (https://gnomad.broadinstitute.org/) and the ones identified in cBioPortal. Patients belonging to two different datasets were considered only one time.

### Quantifications

In cell culture experiments, to quantify the cCas3 signal, the ImageJ software and the Integrated Raw density method were used. We counted four different fields and the integrated density for every field was calculated with an adjusted threshold of 183. t-test was used for statistical analysis. For adhesion tests, three images for each well for each condition were acquired and cells were counted by ImageJ software. Then, the average of each replicate was normalized with the average of control cells for each experiment. Significance was calculated using t Student’s test.

In fly experiments, each experimental point was performed with samples from independent crosses with three replicates per experimental point. Statistical significance was assessed using One-or Two-Way ANOVA, non-parametric (Mann-Whitney) test, Uncorrected Fisher’s LSD test for immunofluorescence data. Survival curve analyses were conducted using the Gehan-Breslow-Wilcoxon and Log-rank (Mantel-Cox) tests. The selected statistical test and sample size are shown in the figure legend below each figure.

For the analysis of larval wing discs acquired at 20× magnification, a selection mask was created around the GFP-positive tissue to quantify surface area, GFP intensity, Cas3, and pH3. The number of positive pixels within the GFP+ area was determined, with surface area expressed in absolute values (pixels) and other measurements reported as the percentage of positive pixels within the GFP+ region.

Quantification of DNA damage-positive cells with γH2Av was conducted manually on a single medial plane chosen in the dorsal portion of the wing pouch, where the cells were all aligned on the same plane. Cas3 quantification and colocalization analysis were conducted manually across the entire Z-stack, as apoptotic cells were sparse and distributed across different planes but remained within the GFP+ area.

For the measurements of tumor-like masses in adult brains, the GFP+ area was measured on a Max Projection of the full Z-stack using FIJI’s “Freehand Selection” tool and quantified with the “Measure” command.

Immunofluorescence analysis of larval brains revealed NBII clusters with complex three-dimensional structures. Neuroblasts (1–8 per hemisphere, anatomically ordered) were identified, and a 10-slice Z-stack (0.2 μm per slice) was generated for each. The GFP+ outer contour was outlined using the “Freehand Selection” tool, creating a ROI mask applied to individual fluorescence channels. The “Threshold” function defined positive signals relative to background, and the occupied area was measured as the percentage of pixels within the ROI. Data obtained from FIJI were visualized as violin plots in Prism, where each point represents a single neuroblast measurement.

## RESULTS

### *STAG2* variants are present in glioblastoma and medulloblastoma patients

To evaluate the presence of *STAG2* variants in patients diagnosed with glioblastoma multiforme or medulloblastoma, we examined the publicly accessible cBioPortal, selecting datasets from 8 studies encompassing 1538 patients. We detected *STAG2* variants in 21 patients (2%), 8 with medulloblastoma and 13 with glioblastoma. These variants were scattered along the *STAG2* gene without any hotspot regions even though 4 out of 8 medulloblastoma samples occurred within a 120bp region. Specifically, 6 missense, 7 nonsense, 8 splicing mutations, and 5 frameshifts. Overall, their CADD score is above the average, suggesting a high impact of these variants on STAG2 function (**Fig. 1A**). Notably, in two patients with medulloblastoma (Group 3 and WNT group), the same mutation, R259*, was reported. Patient details are listed in **suppl. Table 1**.

**Fig. 1.**
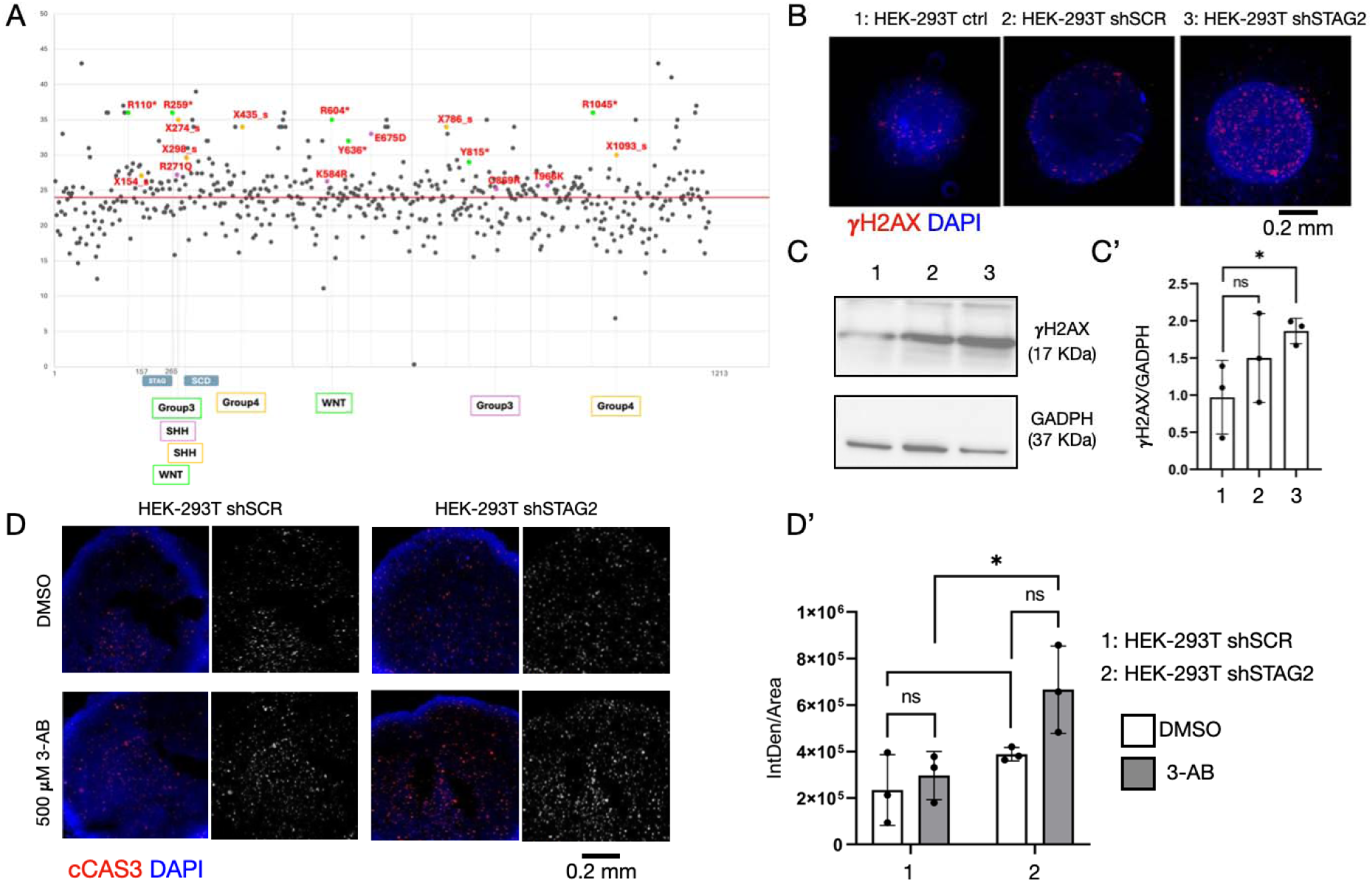
STAG2 variants in brain cancer*, STAG2* depletion and PARP inhibitor treatment in human cells. A Variants in *STAG2* observed in medulloblastoma and glioblastoma. The graph represents the CADD value, a score of variant deleteriousness, for all the variants reported in gnomAD, the genome aggregation database, for *STAG2* (grey) and the variants identified in cBioPortal (in color). Purple represents the missense variants, green represents the nonsense variants and yellow is for the splicing mutations. The frameshift is not reported in the graph. Overall, their CADD score is above the average (red line), suggesting a high impact of these variants on STAG2 functioning. The molecular subtyping of medulloblastoma for the indicated variants is shown below the graph. B Confocal sections of control HEK-293T spheroids or spheroids stably expressing a scrambled control shRNA (shSCR) or a shRNA targeting *STAG2* (shSTAG2) treated to label DNA (DAPI) and DNA damage foci (γH2A.X). C-C’ Western blot analysis of the indicated extracts to detect γ H2A.X and GADPH levels and relative quantification. D-D’ Confocal sections of the indicated spheroids treated as indicated and labeled to detect the DNA (DAPI) and the apoptotic marker cleaved Caspase 3 (cCas3) and relative quantification.

### Synergism of STAG2 knock-down and PARP inhibition in an *in vitro* model

To determine the consequences of reduced *STAG2* activity, we developed an *in vitro* cell model based on STAG2 silencing. To this end, we permanently integrated in HEK-293T cells a lentiviral plasmid expressing a short hairpin against *STAG2* (shSTAG2) or a scrambled sequence (SCR). To assess the knockdown of *STAG2*, we evaluated the expression of both *STAG2* mRNA and protein. In both cases, STAG2 was significantly depleted in HEK-293T shSTAG2, when compared with HEK-293t shSCR and HEK-293T ctrl (**Fig S1A-B**). Depletion did not cause significant changes in cell viability but influenced cell adhesion to the substrate (**Fig S1C-D**).

Considering that the cohesin complex is involved in DNA repair, we next prepared 3D HEK-293T cultures and assessed spheroids for presence of DNA damage. To this end, we analyzed the positivity for γH2A.X, a well-known DNA damage marker, by Western blot analysis. We observed that HEK-293T shSTAG2 spheroids show higher amounts of γH2A.X when compared with HEK-293T shSCR or control spheroids **(Fig 1B-C**).

Given the increased DNA damage in STAG2-depleted cells and the reported susceptibility of cells with aberrant DNA repair mechanisms to PARP inhibitors, we assessed the possible effects of PARP inhibition in our *in vitro* model. Thus, we treated HEK-293T shSTAG2, shSCR and ctrl with 3-Aminobenzamide (3-AB), a potent PARP inhibitor. After 72 hours of treatment, we performed immunofluorescence staining to detect positivity to cleaved caspase 3 (cCas3). We found that HEK-293T shSTAG2 spheroids treated with PARP inhibitor displayed a higher cCas3 positivity than controls treated with vehicle (DMSO; **Fig. 1D**), supporting the possibility of a synthetic lethality induced by combined loss of STAG2 and PARP activity. Overall, these data suggest that the presence of a pharmaco-genetic interaction between STAG2 and PARP activity, that might depend on DNA damage.

### Reduction of *SA1* activity in *Drosophila melanogaster* affects DNA repair *in vivo*

To validate our *in vitro* results, we studied the consequences of somatic depletion of *Drosophila SA1* by expressing *in vivo* TRiP.GL00534 or TRiP.HMS00272, two hairpins to induce RNA interference (RNAi) against *SA1,* in cells of the pouch of larval wing imaginal discs with the GAL4 driver ms1096Bx (*MS>*; **Fig. 2A**). Because TRiP.GL00534 (SA1-RNAi hereafter) resulted in slightly more efficient than TRiP.HMS00272 (**Fig. S2A**), we selected it for further analyses. To achieve *Parp1* depletion, we used an established RNAi line. *Parp1* depletion in epithelial tissue of the wing imaginal disc pouch (*MS>Parp1-RNAi)* caused in nucleolar fragmentation, as previously reported (Boamah *et al*., 2012), indicating efficiently inactivates Parp1 (**Fig. S2B**). Supplementation in the fly food with 3-AB also caused nucleolar fragmentation in epithelial tissue, indicating that the inhibitor is bioactive and was well tolerated at the dosage used for supplementation (**Fig. S2C-D**).

**Fig. 2.**
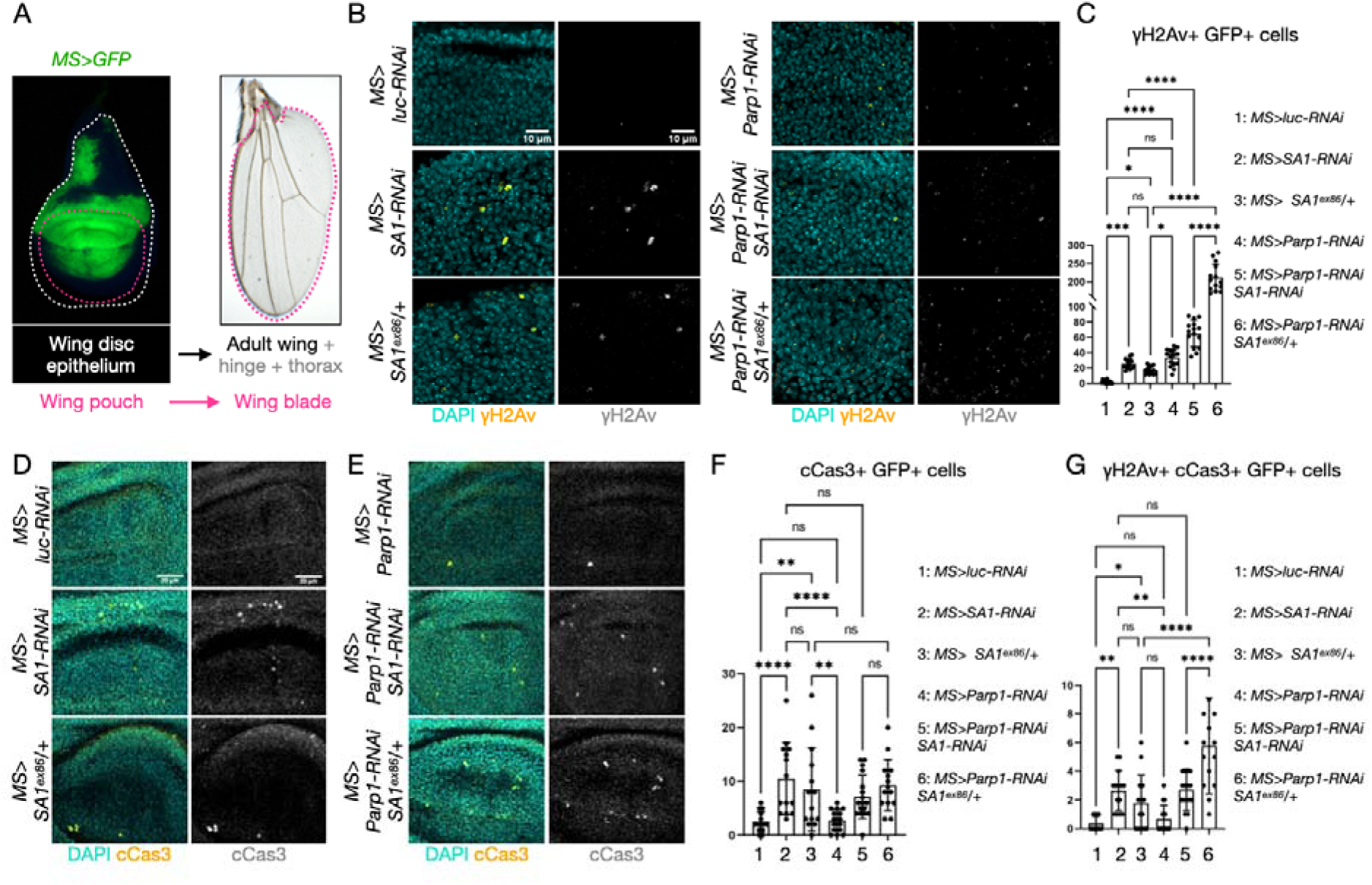
DNA damage and apoptosis upon *in vivo* downmodulation of cohesin and PARP activity. A Schematics of wing imaginal discs illustrating their structure, fate and domain of misexpression using MS>. B-C Confocal sections of the pouch of imaginal discs depleted *in vivo* as indicated, treated to label DNA (DAPI) and DNA damage foci (γH2Av) and relative quantification. D-G Confocal sections of the pouch of imaginal discs depleted *in vivo* as indicated, treated to label cleaved Caspase 3 (cCas3) and relative quantification.

We next immunolocalized DNA damage foci using an antibody against the marker Y2Av in the larval wing imaginal discs, a monolayer epithelial tissue that give rise to the adult wing blade, the hinge and the thorax (**Fig. 2A**). While control *MS>luc-RNAi* cells present no DNA damage foci, we observed that cells of *MS>SA1-RNAi, SA1^ex86^/+* (*SA1* heterozygous)*, or MS>Parp1-RNAi* animals display a significant amount of DNA damage. Interestingly, *Parp1-RNAi* cells that are also depleted or heterozygous for *SA1* display more DNA damage foci, when compared to single manipulations (**Fig. 2B**, quantified in **C**). These data suggest that combined reduction of *SA1* and *Parp1* leads to an additive effect on DNA damage.

To determine whether the observed DNA damage is associated with Caspase-dependent apoptotic cell death, we immuno-localized cCas3. While control *MS>luc-RNAi* or *MS>Parp1-RNAi* cells present only occasional cCas3 staining, we observed that *MS>SA1-RNAi* or *SA1^ex86^/+* cells display a significant amount of Cas3 signal (**Fig. 2D**, quantified in **F**). These data suggest that reduced *SA1* expression but not reduced *Parp1* expression causes Caspase-dependent apoptotic cell death. Consistent with this, when combining reduced *SA1* and *Parp1* expression, no additional Cas3-positive signal is observed *in vivo* (**Fig. 2E**, quantified in **F**). A similar pattern is observed upon quantification of cells that are positive for DNA damage foci and cCas3 expression (**Fig. 2G**).

Together, these data suggest that cells with reduced *SA1* and *Parp1* expression present high level of DNA damage and that a certain proportion of cells with reduced *SA1* expression are likely eliminated by Caspase-dependent apoptosis (Xu *et al*., 2017).

Consistent with the presence of cCas3-positive cells in the larval wing pouch, *MS>SA1-RNAi* adult animals display reduced wings, a sign of uncompensated cell death during tissue development. In contrast, *MS>SA1-RNAi, Parp1* animals display a milder wings reduction (**Fig. 3A**, quantified in **B**). Amelioration of the wing phenotype was also observed upon feeding 3-AB *MS>SA1-RNAi* animals (**Fig. 3C**). Wings appear normal in *MS>Parp1-RNAi* or *SA1^ex86^/+* animals, or in *MS>SA1-RNAi* animals fed 3-AB (**Fig. 3A-B**), indicating that Parp1 inactivation or halving *SA1* dosage do not *per se* affect wing development. These *in vivo* data suggest that concomitant reduction of *SA1* and *Parp1* might sensitize cells towards elimination of the most defective.

**Fig. 3.**
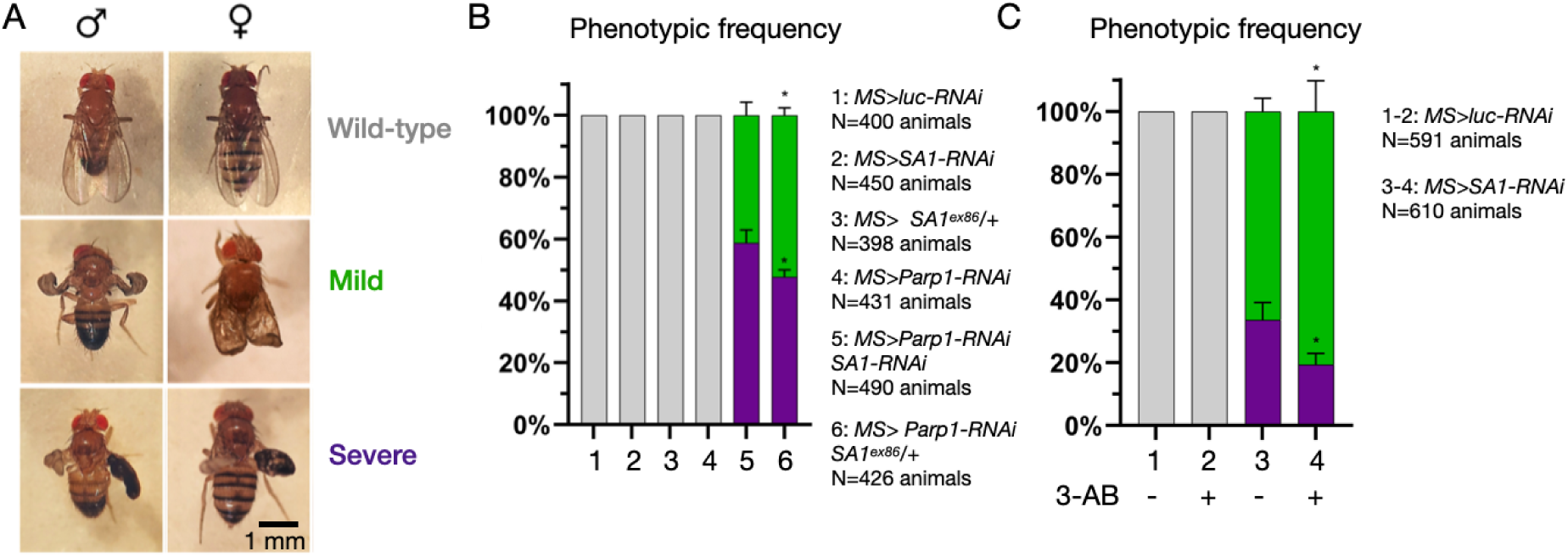
Amelioration of *SA1* depletion phenotypes upon reduction of PARP activity. A Classification of representative wing phenotypes of control animals or upon downmodulation of cohesin and PARP activity. B-C Quantification of the wing phenotypes of animals of the indicated genotypes and treated as indicated. The number of analyzed animals of each condition is indicated in the graph bar. P-value

### Cohesin activity is tumor suppressive during fly brain development

To assess the contribution of cohesin genes to brain tumorigenesis *in vivo*, we first employed a genetic glioma model (Read *et al*., 2009), based on expression in developing larval glial cells of an active form of human EGFR together with expression the PI3K homolog Pi3K92E/Dp110 and of mC8GFP to mark the glial tissue **(Fig. 4A**; *repoGFPEP>* hereafter). Expression in the control glia (*repoGFP>* hereafter) of a mock hairpin targeting *luciferase* (*repoGFP>luc-RNAi*) or depletion of 5 different cohesin genes did not lead to major reduction of glial tissue growth at 120 hours after egg laying (AEL; **Fig. 4B**, quantified in **C**). In contrast, depletion of the cohesin genes *SA1, SMC1* and *SMC3* during gliomagenesis lead to increased glioma growth, as well as to earlier lethality, when compared to the levels of glial overgrowth observed in *repoGFPEP>luc-RNAi* animals (**Fig. 4D**, quantified in **E; Fig. S3**). Overall, these data suggest that some cohesin genes, among which *SA1,* act as tumor suppressors during gliomagenesis *in vivo*.

**Fig. 4.**
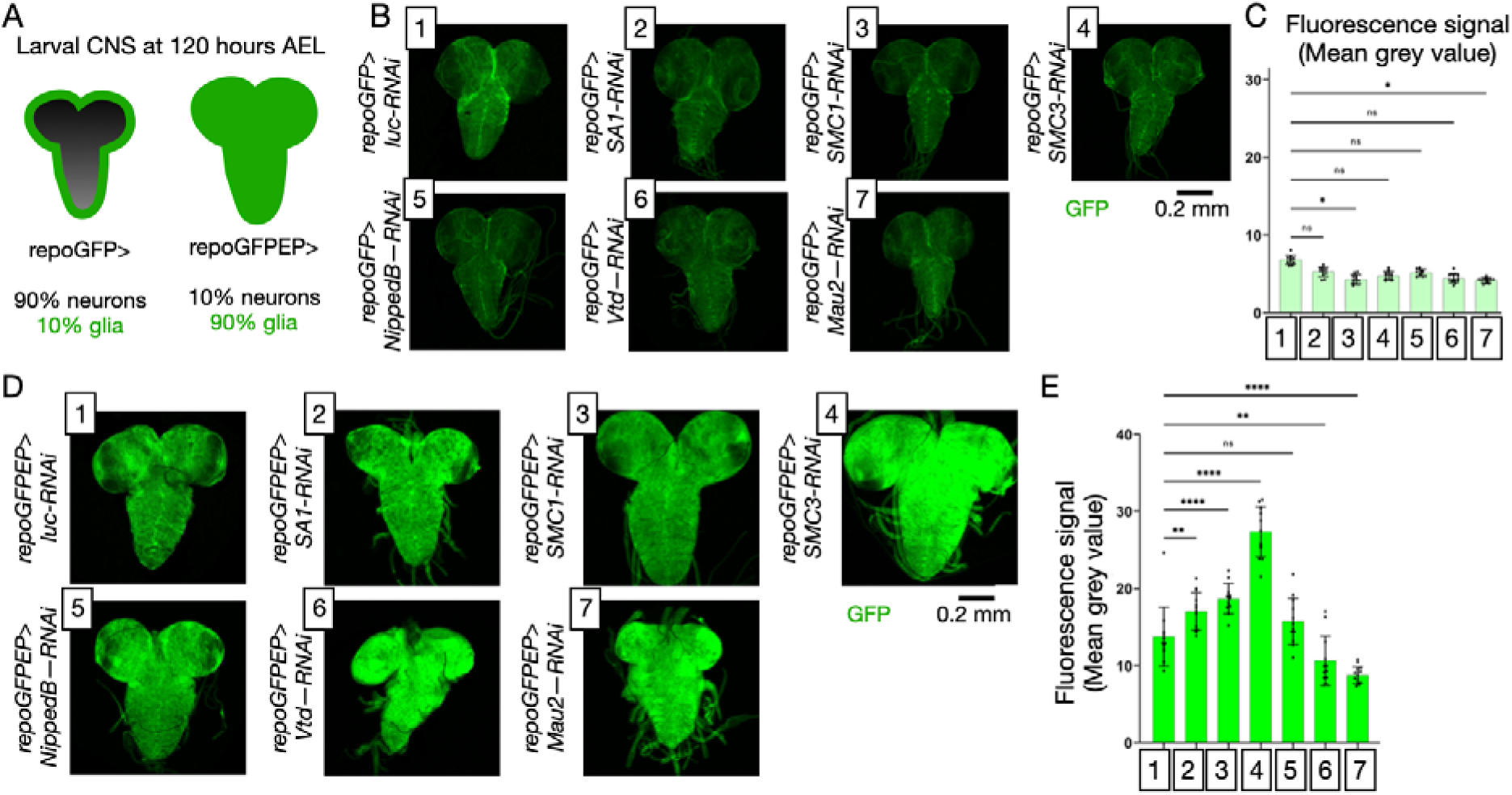
Tumor suppressive activity of cohesin genes in an *in vivo* glioma model. A Schematics of the gliomagenesis model in *Drosophila* larvae. B-E Confocal images of the larval CNS of animals of the indicated genotype and relative quantification.

To further analyze the role of *SA1* in brain tumorigenesis, we next used a genetic background that drives gene expression in larval type II neuroblasts (NBII hereafter; (Neumüller *et al*., 2011)), the 8 neural stem cells (NSC) that generate the neurons and glia of the posterior part of each hemisphere of the larval fly brain (**Fig. 5A**). In contrast to controls, in which NBII drives expression of mCD8::GFP to mark the NBII lineage and of a mock hairpin targeting *luciferase* (*NBII>luc-RNAi*), depletion of the tumor suppressor *brat* (*NBII>brat-RNAi*) led to a massive increase in the number of NBII clusters, while expression of the human brain tumor oncogene *NMYC* (*NBII>NMYC*), or depletion of *SA1 (NBII>SA1-RNAi)*, or the combination of the two manipulations (*NBII>NMYC SA1-RNAi),* do not alter NBII numbers (**Fig. 5B**).

**Fig. 5.**
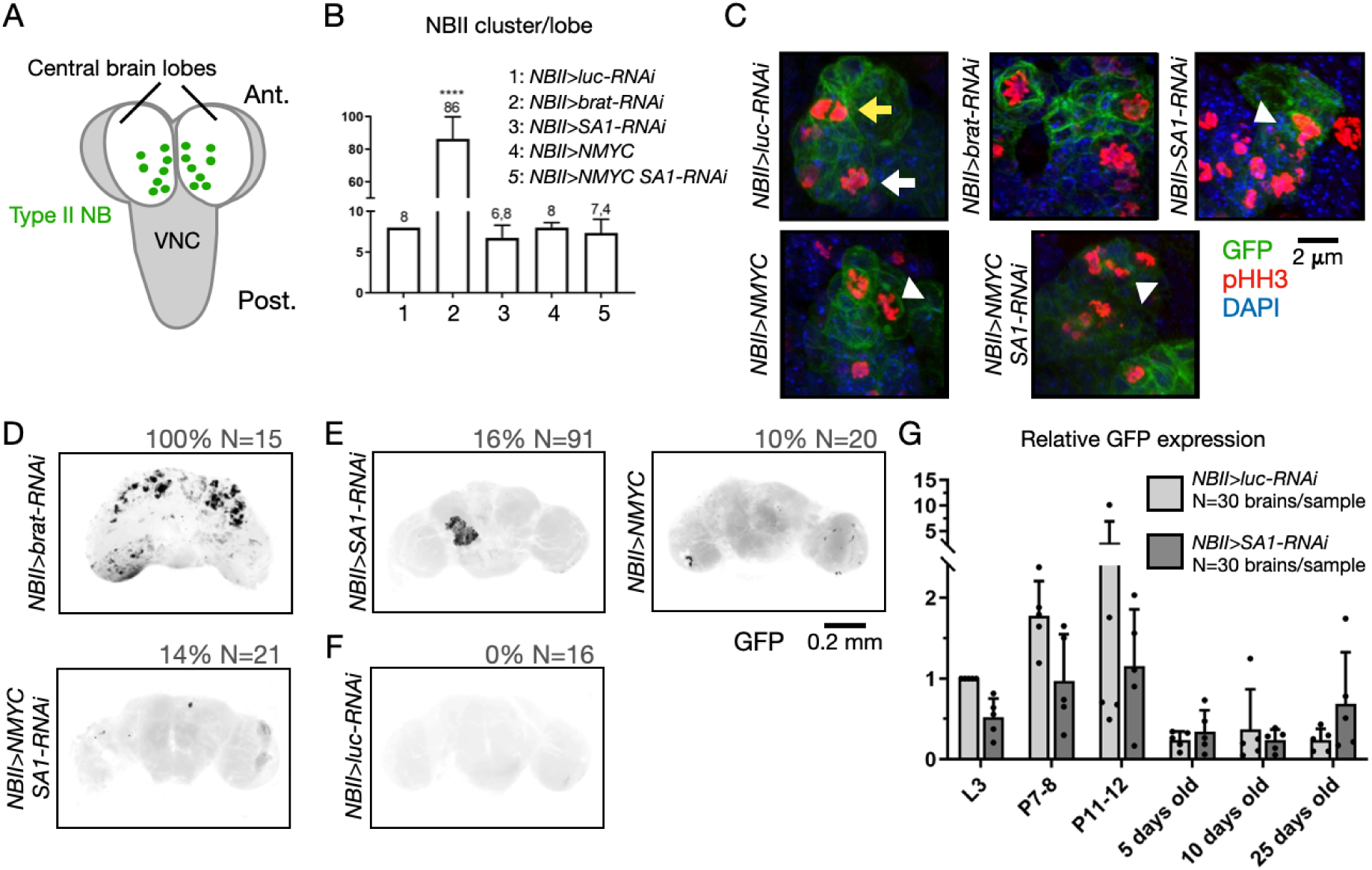
Formation of tumor-like masses upon manipulation of oncogenes and *SA1* expression in neural stem cells. A Schematics of NB positioning in the larval CNS. B Quantification of NBII number upon modulation of the indicated genes. C Representative confocal images of NBII clusters of the indicated genotypes, labeled to detect the indicated cell proliferation marker. The yellow and white arrow point to a normal anaphase and prophase, respectively, while the arrowhead points to examples of aberrant mitotic features. D-F Confocal images of dissected adult brains of the indicated genotype. The percentage of brains with GFP-positive cell masses and the number of brains analyzed (N) is indicated above the panels. G Relative GFP expression by quantitative RT-PCR at distinct developmental times and during adulthood. L3: Third instar larvae; P7-8: white pupae; P11-12: pupae shortly before eclosion.

To visualize dividing cells in NBII cluster, we next immunolabeled with anti-phospo-HistoneH3 (pH3). In *NBII>NMYC*, *NBII>NMYC SA1-RNAi* or *NBII>SA1-RNAi* we observed the formation of aberrant mitotic figures, a defect not observed in control to *NBII>luc-RNAi* or animals or *NBII>brat-RNAi* animal (**Fig. 5C**).

Finally, we wondered whether our manipulations could lead to tumor-like formations in the brain, as previously reported for *brat* depleted animals (Hadjipanayis and Brat, 2017; Reichardt *et al*., 2018). All *NBII>brat-RNAi* animals presented masses of GFP-positive cells nested in the adult brain deriving from NBII (**Fig. 5D**). Interestingly, occasional presence of GFP-positive cells was also observed in *NBII>NMYC, NBII>NMYC SA1-RNAi* or *NBII>SA1-RNAi* animals (**Fig. 5E**), while GFP-positive masses are never recovered in the adult brain of *NBII>luc-RNAi* animals (**Fig. 5F**), as NBII expression is known to abate with full maturation of neuroblast clusters at the end of pupal life. Consistent with this, quantification of mCD8::GFP expression by RT-PCR during development in *NBII>luc-RNAi* controls recapitulates cluster maturation with high expression during larval and pupal stages and background expression in adults. In contrast, *NBII>SA1-RNAi* animals express lower mCD8::GFP levels during larval and pupal life, while they express more mCD8::GFP in aging adults (**Fig. 5G**). These data are consistent with the occasional presence of GFP-positive cells in adult fly brains and suggest that animals with reduced *SA1* expression might display impairment of NBII cluster differentiation. Overall, our results indicate that oncogene expression, cohesin depletion, or both prevent brain tumorigenesis *in vivo*.

### PARP inhibition efficiently improves neuroblast differentiation and lifespan in *SA1* knock-down flies

The emergence of NSC-derived undifferentiated brain cells in *SA1*-depleted adults might result from altered differentiation of larval NBII clusters. To analyze their development, we immunolocalized Miranda (Mira), a marker of the intermediate neural precursors (INP) derived from stem cell, Prospero (Pros), a marker of ganglion mother cells (GMC) that are produced by mature INPs, and Elav, which marks differentiated neurons in L3 larvae (**Fig. 6A**). In control *NBII>luc-RNAi* animals, we observed the expected distribution of Mira-, Pros– and Elav-positive cells emerging from GFP-positive clusters. In sheer contrast, *NBII>SA1-RNAi* clusters accumulate Mira-positive cells at the expense of Pros– and Elav-positive cells. Similar results were obtained in *SA1^ex86^/+* animals (**Fig. 6B**, quantified in **D**). These data suggest that reduced *SA1* expression *per se* might delay or arrest the transition of INP to GMC.

**Fig. 6.**
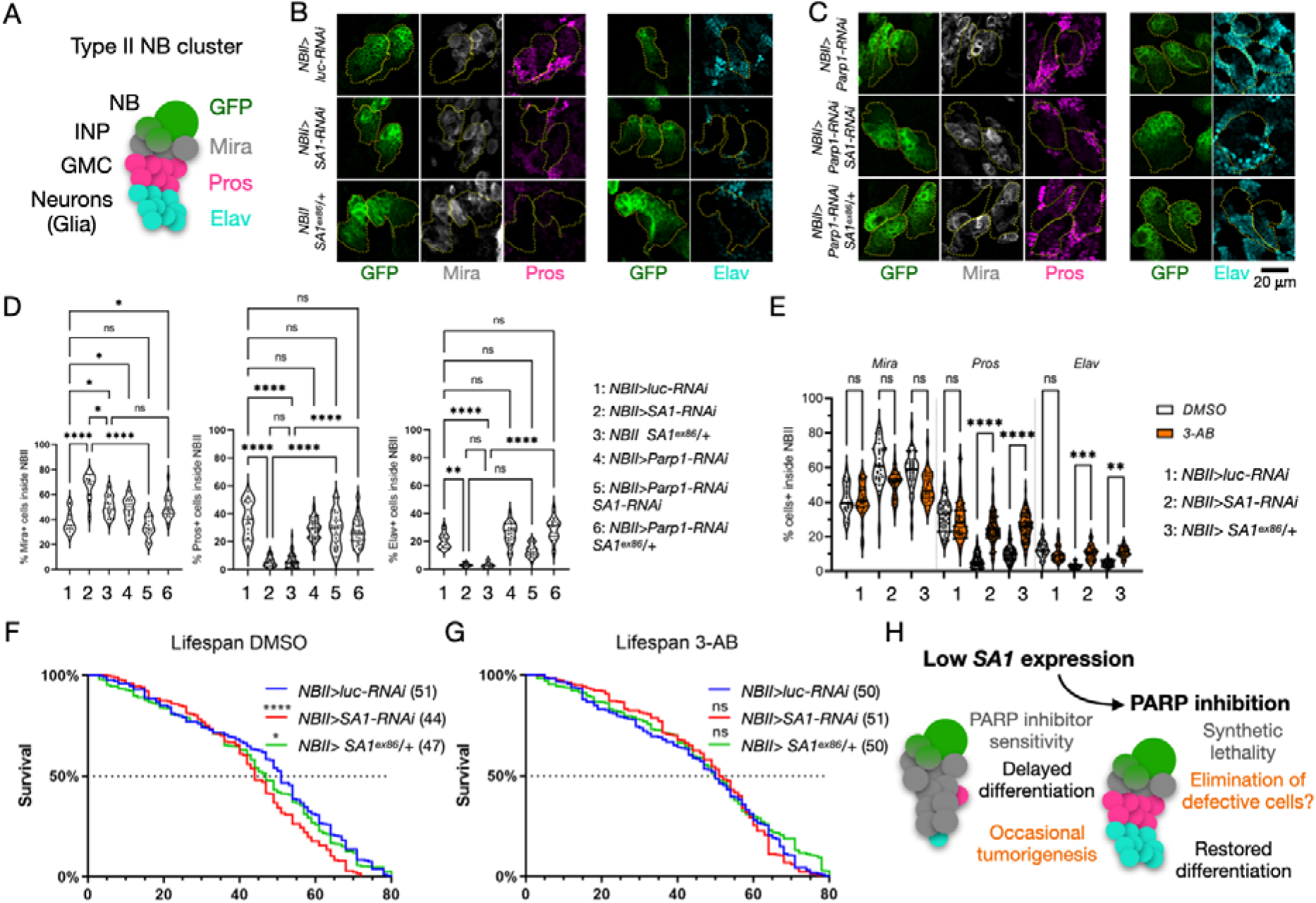
Developmental and lifespan alteration upon *SA1* depletion in developing neuroblasts are rescued by reduction of PARP activity. A Schematics of NBII development and relative markers used to assess it. B-C Representative confocal images of NBII clusters of the indicated genotypes, labeled to detect the indicated differentiation markers. D Quantification of the indicated differentiation marker in NBII cluster of the indicated genotype. E Quantification of the indicated differentiation marker in NBII cluster of the indicated genotype upon supplementation with vehicle (DMSO) alone or 3-AB in vehicle (3-AB). F-G Lifespan of animals of the indicated genotype fed with vehicle alone (DMSO) or 3-AB in vehicle. H A model of brain tumorigenesis based on reduced terminal differentiation.

We next tested whether NBII development is altered by reduction of *Parp1* activity. In *NBII>Parp1-RNAi* neuroblast clusters, we observed a slight increase of Mira-positive cells but no change in Pros– and Elav-positive cells, when compared to *NBII>luc-RNAi* controls (**Fig. 6C**, quantified in **D**), suggesting that decreased Parp1 activity causes only a minor alteration of NBII differentiation. Remarkably, downregulation of *Parp1* ameliorated the developmental delay observed in *SA1* depleted or *SA1^ex86^/+* animals (**Fig. 6C**, quantified in **D**). To determine whether inactivation of Parp1 during NBII differentiation could also be achieved pharmacologically, we fed animals with 3-AB. Consistent with our genetic data, we observed that 3-AB supplementation *per se* does not affect NBII development. However, it partially phenocopies the effect of *Parp1* depletion, with amelioration of Pros– and Elav-positive cell differentiation (**Fig. 6E**).

Despite the overall morphology of the adult brain of control, *SA1*-downregulated and *SA1* heterozygous animals appears unaffected (**Fig. S4**), we observed that compared to *NBII>luc-RNAi*, *NBII>SA1-RNAi* animals display a 17% reduction in median survival. Similar data were obtained in control animals heterozygous for a null *SA1* allele (*NBII SA1^ex86^/+;* **Fig. 6F**). Consistent with the effect on NBII differentiation, while 3-AB supplementation did not significantly alter the lifespan of control *NBII>luc-RNAi* animals, it improved the lifespan of animals with reduced *SA1* expression (**Fig. 6G**).

Overall, these results suggest that SA1 supports neuroblast cluster development and that the defects observed upon *SA1* reduction are rescued by impairment of PARP activity.

## DISCUSSION

Database analyses highlighted a possible role of STAG2 in brain tumors, particularly in glioblastoma and medulloblastoma. As reported variants are predicted to result in STAG2 haploinsufficiency, we exploited an *in vitro* and *in vivo* system for modelling the contribution of reduced *STAG2* activity to relevant processes. Using 293T cells with stable downregulation as a 3D *in vitro* model, we observed increased DNA damage. Interestingly, exposing depleted cells to PARP inhibitors we observed high levels of cell death suggesting the possibility of synthetic lethal interaction between reduced *STAG2* and PARP activity. Our *in vivo* data based on depletion or heterozygosity of *SA1,* the fly homolog of *STAG2* confirm that cohesin genes act as tumor suppressors. The also show that synthetic lethal interaction are occurring *in vivo*. Because cohesin components play multiple cellular roles, our data, do not clarify which of them is tumor suppressive.

While we cannot conclude that accumulation of DNA damage upon reduction of *SA1* expression in *Drosophila* contribute to tumorigenesis, such phenotypes are reminiscent of those reported in response to replication stress upon cohesin removal, or PARP inhibition, or oncogenic MYC activity (Benedict et al. 2020; Colicchia et al. 2017; Peripolli et al. 2024). Consistent with this, we obtained brain tumor-like masses in flies by NSC-directed overexpression of either NMYC, by *SA1* depletion or both. PARP inhibition *per se* is sufficient to cause accumulation of DNA damage and the combination of PARP inhibition and reduction of *SA1* expression results in additive effects. Our results are in line with the reports that glioblastoma cells with *STAG2* mutations show increase in DNA damage markers and cell cycle arrest caused by replication stress when treated with PARP inhibitors (McLellan et al. 2012; Tothova et al. 2013; Bailey et al. 2020) and with a clinical trial exploring PARP inhibition in blood cancers with mutations in cohesin genes (https://clinicaltrials.gov/study/NCT03974217), and suggest that a synthetic lethal interaction that PARP inhibition in medulloblastoma *in vitro* models (Price and Lau 2023) might involve cohesin activity. The additive effects that we observe *in vivo* correlate with phenotypic amelioration in two different organs and improve lifespan of the animals. Thus, our fly and human cell genetic backgrounds are likely to be informative to dissect the impact of cohesin activity on synthetic lethality induced by PARP inhibition in the context of brain tumorigenesis.

Importantly, a recent study in *Drosophila* has found that the sole alteration of epigenetic regulation of chromatin by impairing Polycomb activity is sufficient to promote tumorigenesis in absence of driver mutations (Parreno et al. 2024). Thus, regulation of chromatin architecture could be the main function of cohesin proteins relevant to tumor suppression. Despite this, it is likely that the contribution of reduced *SA1* expression to brain tumorigenesis might be pleiotropic, considering that in our experimental systems upon *SA1* downregulation we also observe defective mitotic figures and accumulation of DNA damage.

Our data showed that correct *SA1* expression supports neuroblast differentiation during larval life and prevents the persistence of NBII-derived INPs, few of which might correspond to the NBII-derived cells recovered in adult brains. Cohesin genes have been implicated in axonal pruning (Schuldiner et al. 2008), suggesting that elimination of persistent INP generated by low *SA1* expression might fail post mitotically. It has also been shown that precise control of NB cell elimination during embryonic nervous system development depends on *SA1* activity downstream of regulation of the H3K27me3 state of chromatin by Notch (Arya et al. 2019). Remarkably, epigenetic regulation by Polycomb proteins increases the activity of stemness genes during asymmetric cell division of NBII by elevating H3K27me3 levels at cis-regulatory elements in INP cells. These authors suggest that a failure of this process could reduce Notch activity and thereby promote INP proliferation instead of maintaining their stemness (Rajan et al. 2023). In this context, Notch activity is also known to be repressed by the tumor suppressor Brat (human TRIM3) to promote differentiation of immature neural precursors (Hadjipanayis and Brat 2017). Consistent with this, the tumor-like masses observed in adult brain of flies with reduced NSC expression of *SA1*, resemble those more abundantly obtained upon *brat* depletion. Taken together, our evidence suggests that *SA1* reduction might alter Notch regulation of NBII development toward proliferation of INPs and away from their elimination at the correct developmental times, eventually leading to tumorigenesis (**Fig. 6H**).

Reduced cohesin activity might impact also other tumor-relevant cell behaviors. In fact, *SA1* depletion has been recently shown to increase migration of fly tumor cells of epithelial origin (Canales Coutiño et al. 2020). Thus, it will be interesting to study also whether the brain tumor-like masses observed in our experiments upon *SA1* depletion have migrated away from sites of NBII development. Interestingly, in a *in vivo* gliomagenesis model that highlights proliferative effects, *SA1, SMC1,* and *SMC3* depletion all promoted tissue growth, an effect opposite to that corresponding depletion in control animals. These data indicate that cohesin genes might play tumor suppressive roles also after differentiation of glial cells perhaps relevant to glioblastoma, a tumor in which we and others have reported inactivating *STAG2* mutations. However, how reduced cohesin genes expression might promote tissue growth remains to be determined.

## AUTHOR CONTRIBUTIONS STATEMENT

TV and VM designed the study. ST conducted all the *Drosophila* experiments with the technical help of FL under the supervision of TV. AL and SV performed the experiments in human cells and the computational analysis under the supervision of CG and VM. TV and VM wrote the manuscript with the help of ST and AL.

## DATA AND RESOURCE AVAILABILITY

All relevant data and details of resources can be found within the article and its supplementary information.

## ACKNOWLEDGEMENTS

The authors are thankful of the support of the microscopy facility NOLIMITS of the University of Milan and the Bloomington and Vienna *Drosophila* Stock center. We are grateful to Cedric Maurange (Aix Marseille University) and Kim McKim (Rutgers University) for providing fly stocks.

## Supplementary figures

**Fig. S1.**
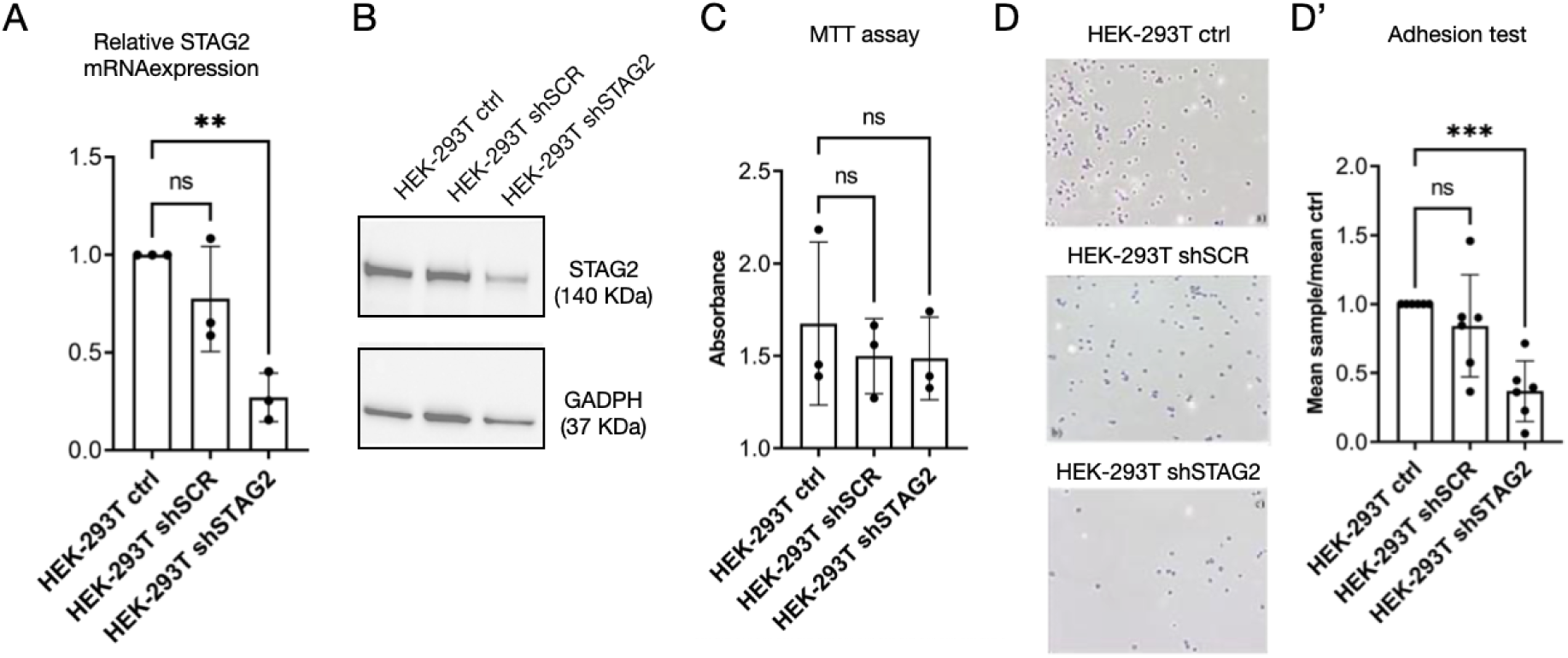
*STAG2* depletion affects cell adhesion. A RT-PCR quantification of *STAG2* mRNA expression in control cells or cells expressing the indicated shRNA. B Western blot analysis of STAG2 and GADPH expression in control cells or cells expressing the indicated hairpin. The blot of GADPH to normalize is the same of Fig. 1C. C Determination of cell density of control cells or cells expressing the indicated hairpin D Substrate adhesion of control cells or cells expressing the indicated shRNA and relative quantification.

**Fig. S2.**
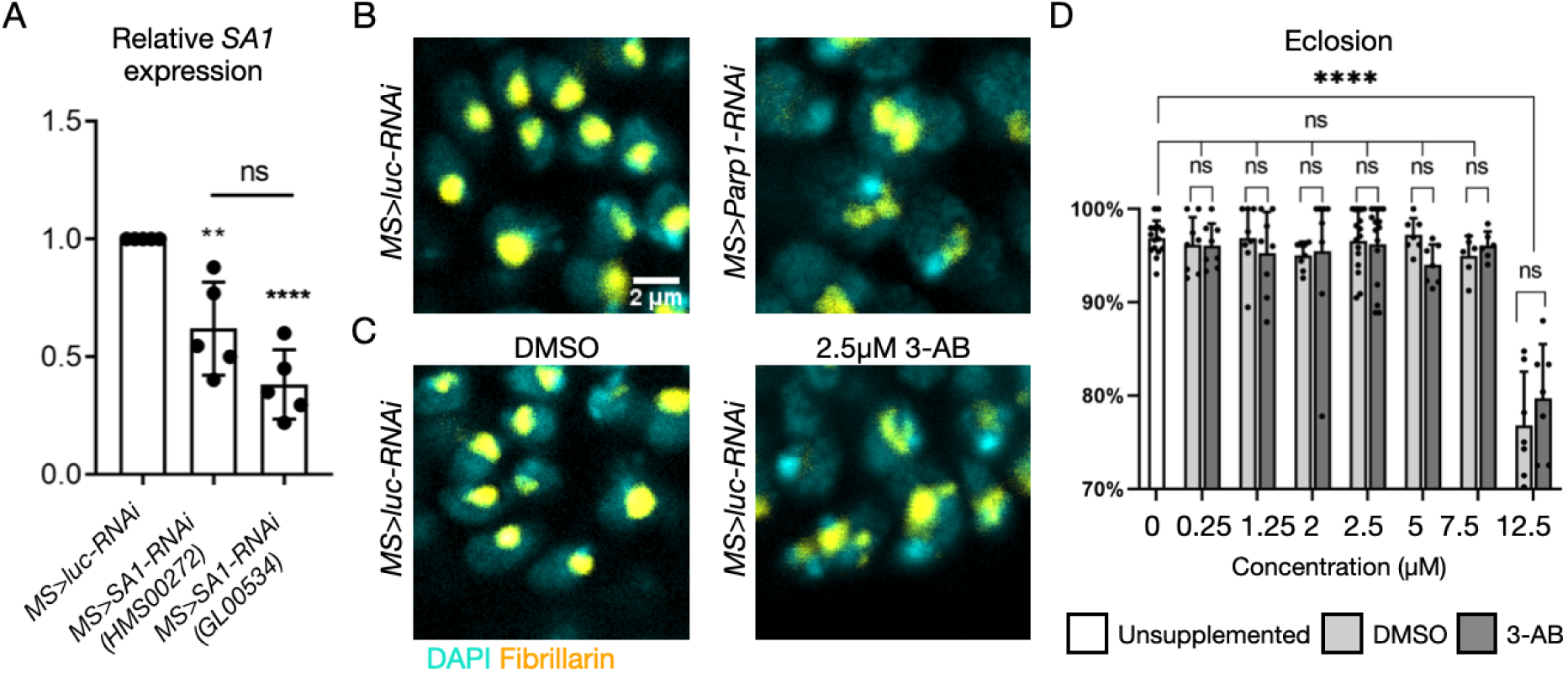
*SA1* downregulation and reduction of PARP activity. A RT-PCR quantification of *SA1* mRNA expression in extracts of wing imaginal discs of the indicated genotypes. B-C High magnification confocal images of nuclei of the wing pouch of animals of the indicated genotypes treated as indicated that have been labeled to detect the DNA (DAPI) and the nucleolar marker fibrillarin. D Eclosion rates of control animals untreated or fed with the indicated concentration of vehicle alone (DMSO) or 3-AB in vehicle (3-AB).

**Fig. S3.**
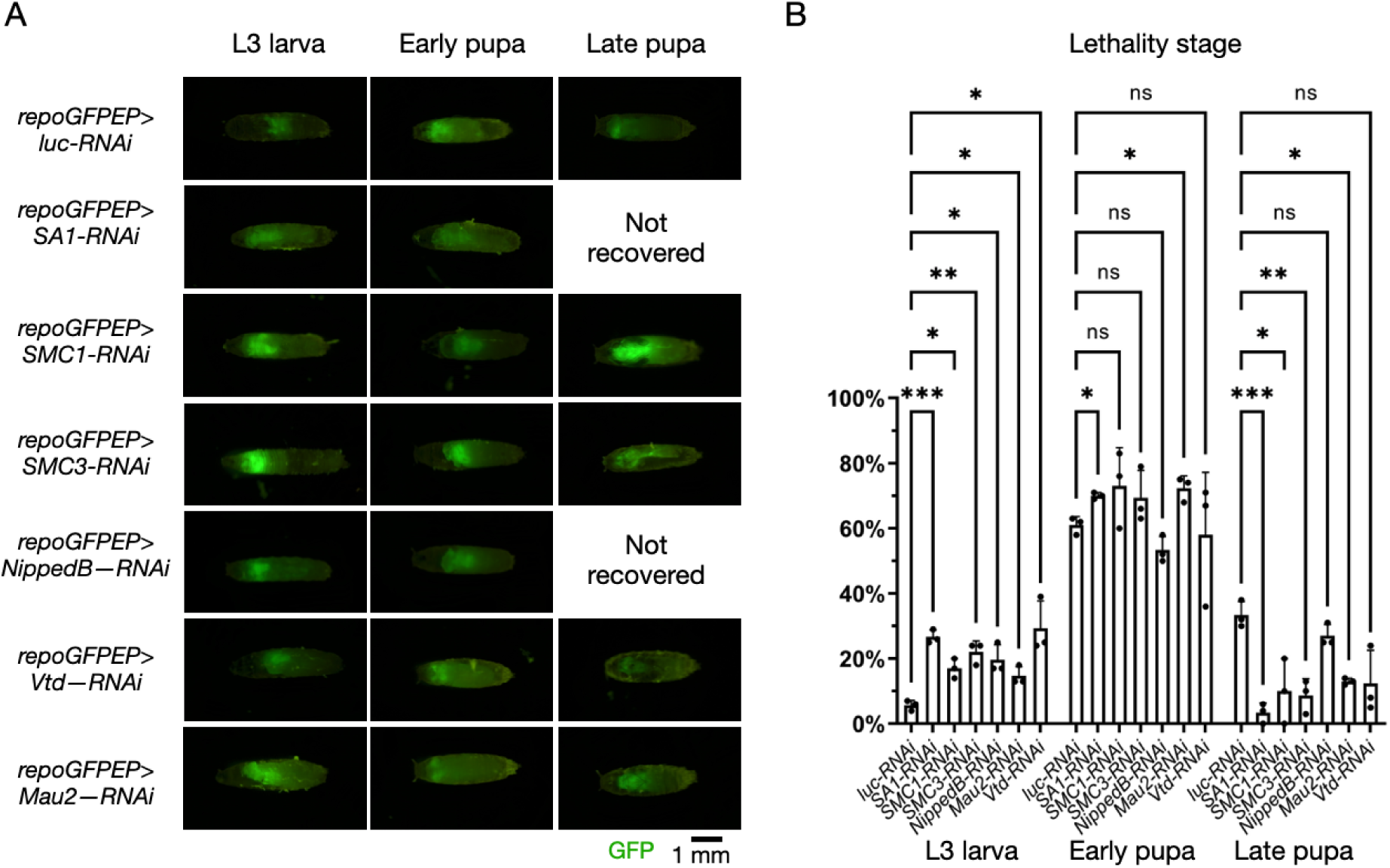
Lethality stage of animals with reduced cohesin gene expression. A-B Representative confocal images of *repo>GFPEP* larve depleted of the indicated cohesin gene and quantification of their lethality stage.

**Fig. S4.**
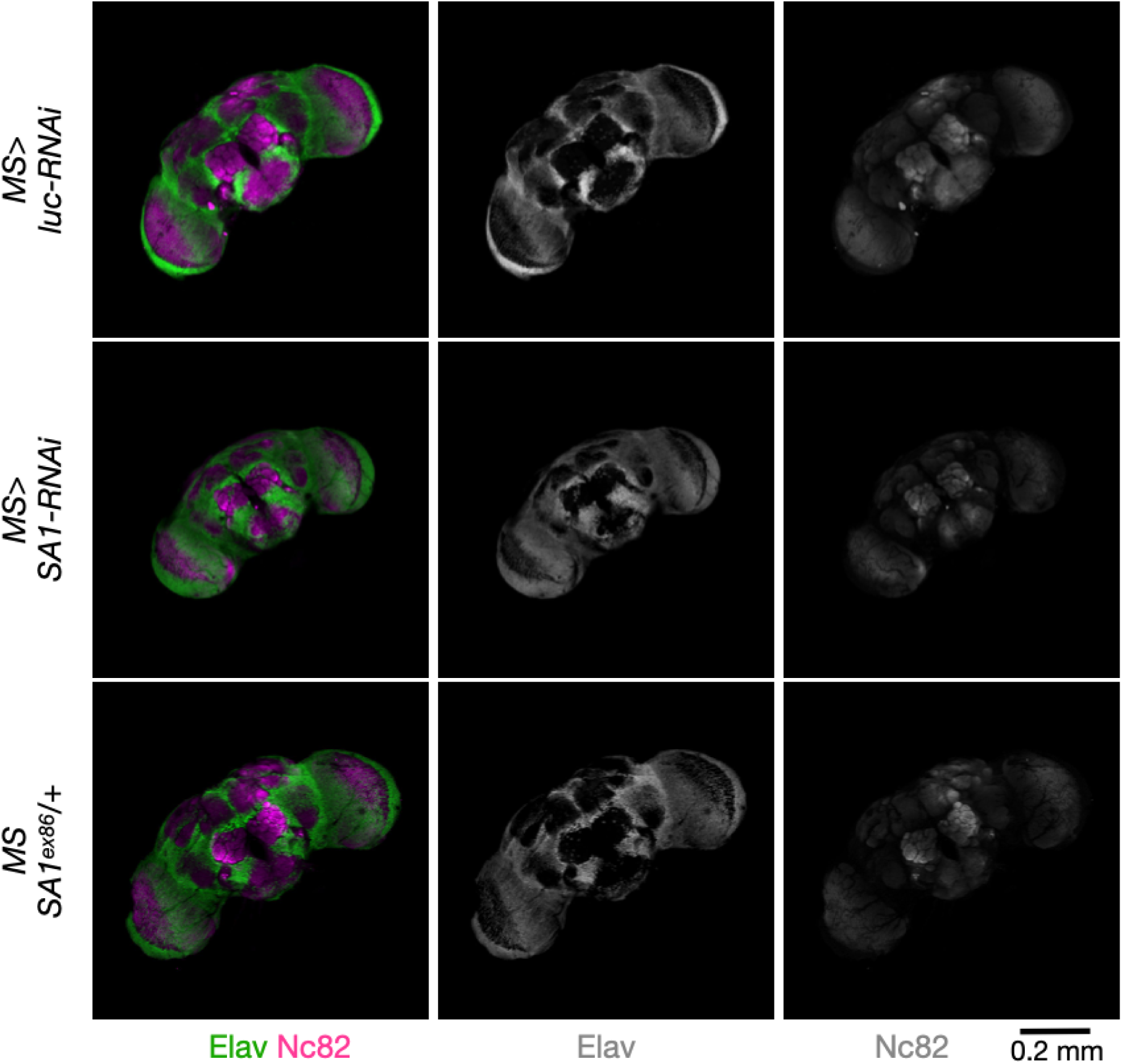
Brain morphology with differing levels of SA1 expression. Confocal images of dissected adult brains of the indicated genotype, immunolabeled to detect the indicated protein markers.

## REFERENCES

1. Arruda, N. L. et al. (2020) ‘Distinct and overlapping roles of STAG1 and STAG2 in cohesin localization and gene expression in embryonic stem cells’, Epigenetics and Chromatin, 13(1). doi: 10.1186/s13072-020-00353-9.

2. Arya, R. et al. (2019) ‘A Cut/cohesin axis alters the chromatin landscape to facilitate neuroblast death’, Development (Cambridge), 146(9). doi: 10.1242/dev.166603.

3. Bailey, M. et al. (2013) ‘Glioblastoma Cells Containing Mutations in the Cohesin Component STAG2 Are Sensitive to PARP Inhibition’ Molecular Cancer Therapeutics, 13, 724–732. doi.10.1158/1535-7163.MCT-13-0749.

4. Balbás-Martínez, C. et al. (2013) ‘Recurrent inactivation of STAG2 in bladder cancer is not associated with aneuploidy’, Nature Genetics, 45(12), pp. 1464–1469. doi: 10.1038/ng.2799.

5. Bayés, M. et al. (2001) ‘Evaluation of the Stag3 gene and the synaptonemal complex in a rat model (as/as) for male infertility’, Molecular Reproduction and Development, 60(3). doi: 10.1002/mrd.1104.

6. Benedict, B. et al. (2020) ‘WAPL-Dependent Repair of Damaged DNA Replication Forks Underlies Oncogene-Induced Loss of Sister Chromatid Cohesion’, Developmental Cell, 52(6). doi: 10.1016/j.devcel.2020.01.024.

7. Boamah, E. K. et al. (2012) ‘Poly(ADP-ribose) polymerase 1 (PARP-1) regulates ribosomal biogenesis in Drosophila nucleoli’, PLoS Genetics, 8(1), pp. 1–14. doi: 10.1371/journal.pgen.1002442.

8. Brennan, C. W. et al. (2013) ‘The somatic genomic landscape of glioblastoma’, Cell, 155(2). doi: 10.1016/j.cell.2013.09.034.

9. Brohl, A. S. et al. (2014) ‘The Genomic Landscape of the Ewing Sarcoma Family of Tumors Reveals Recurrent STAG2 Mutation’, PLoS Genetics, 10(7). doi: 10.1371/journal.pgen.1004475.

10. Canales Coutiño, B., et al. (2020) ‘A Genetic Analysis of Tumor Progression in Drosophila Identifies the Cohesin Complex as a Suppressor of Individual and Collective Cell Invasion’, iScience, 23(6), p. 101237. doi: 10.1016/j.isci.2020.101237.

11. Colicchia, V. et al. (2017) ‘PARP inhibitors enhance replication stress and cause mitotic catastrophe in MYCN-dependent neuroblastoma’, Oncogene, 36(33), pp. 4682–4691. doi: 10.1038/onc.2017.40.

12. Crompton, B. D. et al. (2014) ‘The genomic landscape of pediatric Ewing sarcoma’, Cancer Discovery, 4(11). doi: 10.1158/2159-8290.CD-13-1037.

13. D’Andrea, A. D. (2018) ‘Mechanisms of PARP inhibitor sensitivity and resistance’, DNA Repair, 71, pp. 172–176. doi: 10.1016/j.dnarep.2018.08.021.

14. Gause, M. et al. (2008) ‘Functional links between Drosophila Nipped-B and cohesin in somatic and meiotic cells’, Chromosoma, 117(1), pp. 51–66. doi: 10.1007/s00412-007-0125-5.

15. Guo, G. et al. (2013) ‘Whole-genome and whole-exome sequencing of bladder cancer identifies frequent alterations in genes involved in sister chromatid cohesion and segregation’, Nature Genetics, 45(12). doi: 10.1038/ng.2798.

16. Hadjipanayis, C. and Brat, D. J. (2017) ‘Drosophila Brat and human ortholog TRIM3 maintain stem cell equilibrium and suppress brain tumorigenesis by attenuating Notch nuclear transport’, Cancer Res, 76(8), pp. 2443–2452. doi: 10.1158/0008-5472.CAN-15-2299.Drosophila.

17. Hill, V. K., Kim, J. S. and Waldman, T. (2016) ‘Cohesin mutations in human cancer’, Biochimica et Biophysica Acta – Reviews on Cancer. doi: 10.1016/j.bbcan.2016.05.002.

18. Jones, D. T. W. et al. (2012) ‘Dissecting the genomic complexity underlying medulloblastoma’, Nature, 488(7409). doi: 10.1038/nature11284.

19. Kline, A. D. et al. (2018) ‘Diagnosis and management of Cornelia de Lange syndrome: first international consensus statement’, Nature Reviews Genetics. doi: 10.1038/s41576-018-0031-0.

20. Kon, A. et al. (2013) ‘Recurrent mutations in multiple components of the cohesin complex in myeloid neoplasms’, Nature Genetics, 45(10). doi: 10.1038/ng.2731.

21. De Koninck, M. and Losada, A. (2016) ‘Cohesin mutations in cancer’, Cold Spring Harbor Perspectives in Medicine, 6(12). doi: 10.1101/cshperspect.a026476.

22. Luo, J. et al. (2024) ‘An update on small molecule compounds targeting synthetic lethality for cancer therapy’, European Journal of Medicinal Chemistry, 278, p. 116804. doi: 10.1016/J.EJMECH.2024.116804.

23. Maurange, C. (2020) ‘Temporal patterning in neural progenitors: From Drosophila development to childhood cancers’, DMM Disease Models and Mechanisms, 13(7). doi: 10.1242/dmm.044883.

24. McLellan, J. et al. (2012) ‘Synthetic Lethality of Cohesins with PARPs and Replication Fork Mediators’ PLoS Genetics, 8. doi.10.1371/journal.pgen.1002574.

25. Mekonnen, N., Yang, H. and Shin, Y. K. (2022) ‘Homologous Recombination Deficiency in Ovarian, Breast, Colorectal, Pancreatic, Non-Small Cell Lung and Prostate Cancers, and the Mechanisms of Resistance to PARP Inhibitors’, Frontiers in Oncology, 12. doi: 10.3389/fonc.2022.880643.

26. Mondal, G. et al. (2019) ‘A requirement for STAG2 in replication fork progression creates a targetable synthetic lethality in cohesin-mutant cancers’, Nature Communications, 10(1). doi: 10.1038/s41467-019-09659-z.

27. Di Nardo, M., Pallotta, M. M. and Musio, A. (2022) ‘The multifaceted roles of cohesin in cancer’, Journal of Experimental and Clinical Cancer Research. doi: 10.1186/s13046-022-02321-5.

28. Nasmyth, K. (2011) ‘Cohesin: A catenase with separate entry and exit gates?’, Nature Cell Biology. doi: 10.1038/ncb2349.

29. Nasmyth, K. and Haering, C. H. (2009) ‘Cohesin: Its roles and mechanisms’, Annual Review of Genetics, 43, pp. 525–558. doi: 10.1146/annurev-genet-102108-134233.

30. Neumüller, R. A. et al. (2011) ‘Genome-wide analysis of self-renewal in Drosophila neural stem cells by transgenic RNAi’, Cell Stem Cell, 8(5), pp. 580–593. doi: 10.1016/j.stem.2011.02.022.

31. Northcott, P. A. et al. (2017) ‘The whole-genome landscape of medulloblastoma subtypes’, Nature, 547(7663), pp. 311–317. doi: 10.1038/nature22973.

32. Ostrovsky, A., Cachero, S. and Jefferis, G. (2013) ‘Clonal analysis of olfaction in Drosophila: Generation of flies with mosaic labeling’, Cold Spring Harbor Protocols, 8(4). doi: 10.1101/pdb.prot071712.

33. Paglia, S. et al. (2017) ‘Failure of the PTEN/aPKC/Lgl axis primes formation of adult brain tumours in drosophila’, BioMed Research International, 2017. doi: 10.1155/2017/2690187.

34. Parreno, V. et al. (2024) ‘Transient loss of Polycomb components induces an epigenetic cancer fate’, Nature, 629(8012). doi: 10.1038/s41586-024-07328-w.

35. Peripolli, S. et al. (2024) ‘Oncogenic c-Myc induces replication stress by increasing cohesins chromatin occupancy in a CTCF-dependent manner’, Nature Communications, 15(1). doi: 10.1038/s41467-024-45955-z.

36. Phan, A. et al. (2019) ‘Stromalin Constrains Memory Acquisition by Developmentally Limiting Synaptic Vesicle Pool Size’, Neuron, 101(1), pp. 103–118.e5. doi: 10.1016/j.neuron.2018.11.003.

37. Poeran (2016) ‘Divergent clonal selection dominates medulloblastoma at recurrence’, Nature, 176(12), pp. 139–148. doi: doi:10.1038/nature16478. Divergent.

38. Price, L. and Lau, B. (2023) ‘Mdb-36. sonidegib, gant61, and veliparib treatment both alone and in combination increases radiation sensitivity in pediatric sonic hedgehog medulloblastoma’, Neuro-Oncology, 25(Supplement_1), pp. i70–i70. doi: 10.1093/neuonc/noad073.268.

39. Pugh, T. J. et al. (2012) ‘Medulloblastoma exome sequencing uncovers subtype-specific somatic mutations’, Nature, pp. 106–110. doi: 10.1038/nature11329.

40. Rajan, A. et al. (2023) ‘Low-level repressive histone marks fine-tune gene transcription in neural stem cells’, eLife, 12. doi: 10.7554/eLife.86127.

41. Read, R. D. et al. (2009) ‘A Drosophila model for EGFR-Ras and PI3K-dependent human glioma’, PLoS Genetics, 5(2). doi: 10.1371/journal.pgen.1000374.

42. Reichardt, I. et al. (2018) ‘The tumor suppressor Brat controls neuronal stem cell lineages by inhibiting Deadpan and Zelda’, EMBO reports, 19(1), pp. 102–117. doi: 10.15252/embr.201744188.

43. Robinson, G. et al. (2012) ‘Novel mutations target distinct subgroups of medulloblastoma’, Nature, 488(7409). doi: 10.1038/nature11213.

44. Rollins, R. A. et al. (2004) ‘ Drosophila Nipped-B Protein Supports Sister Chromatid Cohesion and Opposes the Stromalin/Scc3 Cohesion Factor To Facilitate Long-Range Activation of the cut Gene’, Molecular and Cellular Biology, 24(8). doi: 10.1128/mcb.24.8.3100-3111.2004.

45. Romero-Pérez, L. et al. (2019) ‘STAG Mutations in Cancer’, Trends in Cancer, 5(8), pp. 506–520. doi: 10.1016/j.trecan.2019.07.001.

46. Schuldiner, O. et al. (2008) ‘piggyBac-Based Mosaic Screen Identifies a Postmitotic Function for Cohesin in Regulating Developmental Axon Pruning’, Developmental Cell, 14(2). doi: 10.1016/j.devcel.2007.11.001.

47. Solomon, D. A. et al. (2011) ‘Mutational Inactivation of STAG2 Causes Aneuploidy in Human Cancer’, Science, 333(6045), pp. 1039–1043. doi: 10.1126/science.1203619.Mutational.

48. Solomon, D. A. et al. (2013) ‘Frequent truncating mutations of STAG2 in bladder cancer’, Nature Genetics, 45(12). doi: 10.1038/ng.2800.

49. Taylor, C. F. et al. (2014) ‘Frequent inactivating mutations of STAG2 in bladder cancer are associated with low tumour grade and stage and inversely related to chromosomal copy number changes’, Human Molecular Genetics, 23(8). doi: 10.1093/hmg/ddt589.

50. Thomas, S. E. et al. (2005) ‘Identification of Two Proteins Required for Conjunction and Regular Segregation of Achiasmate Homologs in Drosophila Male Meiosis’, Cell, 123(4), pp. 555–568. doi: 10.1016/j.cell.2005.08.043.

51. Thota, S. et al. (2014) ‘Genetic alterations of the cohesin complex genes in myeloid malignancies.’, Blood, 124(11). doi: 10.1182/blood-2014-04-567057.

52. Tirode, F. et al. (2014) ‘Genomic landscape of ewing sarcoma defines an aggressive subtype with co-association of STAG2 and TP53 mutations’, Cancer Discovery, 4(11). doi: 10.1158/2159-8290.CD-14-0622.

53. Tothova, Z. et al. (2020) ‘Cohesin mutations alter DNA damage repair and chromatin structure and create therapeutic vulnerabilities in MDS/AML’ JCI Insight, 6. doi.10.1172/jci.insight.142149.

54. Wang, L. B. et al. (2021) ‘Proteogenomic and metabolomic characterization of human glioblastoma’, Cancer Cell, 39(4). doi: 10.1016/j.ccell.2021.01.006.

55. Weinstein, J. N. et al. (2014) ‘Comprehensive molecular characterization of urothelial bladder carcinoma’, Nature, 507(7492). doi: 10.1038/nature12965.

56. Wu, J. S. and Luo, L. (2006) ‘A protocol for dissecting Drosophila melanogaster brains for live imaging or immunostaining’, Nature Protocols, 1(4). doi: 10.1038/nprot.2006.336.

57. Xu, P. et al. (2017) ‘Flavonoids of Rosa roxburghii Tratt Exhibit Anti-Apoptosis Properties by Regulating PARP-1/AIF’, Journal of Cellular Biochemistry, 118(11). doi: 10.1002/jcb.26049.

58. Zhao, J. et al. (2019) ‘Immune and genomic correlates of response to anti-PD-1 immunotherapy in glioblastoma’, Nature Medicine, 25(3). doi: 10.1038/s41591-019-0349-y.

